# Evaluation of annual ryegrass genotypes for forage yield and blast resistance

**DOI:** 10.64898/2025.12.18.695135

**Authors:** Dediel Rocha, Luiz Hernane Favero, Joseli Stradiotto Neto, Ulisses Arruda Cordova

**Affiliations:** Researcher at Epagri, Lages,SC, CEP 88502-970; Studenr of Agronomy CAV/Udesc, Lages,SC

**Keywords:** *Lolium multiflorum*, *Pyricularia oryzae*, genotype x environment interaction (GxA)

## Abstract

The annual ryegrass (*Lolium multiflorum* Lam.) is a key winter forage, but its productivity is limited by diseases, especially blast, caused by *Magnaporthe oryzae* (syn. *Pyricularia oryzae*). This study aimed to evaluate the productive performance and reaction to blast of commercial cultivars and advanced lines of annual ryegrass in two edaphoclimatic environments in Santa Catarina, Brazil: Lages (Cfb climate) and Agronômica (Cfa climate). The experiment utilized a randomized block design with eight genotypes and four replicates. Variables analyzed included dry matter yield and blast severity. A bifactorial analysis of variance (genotype × local) and Spearman correlation were performed. The results revealed a significant genotype × environment (GxA) interaction for both dry matter yield and blast severity, indicating a dependence on local conditions. Blast pressure was drastically higher in Agronômica, which is climatically more favorable to the disease. Annual ryegrass genotypes ‘Altovale’ and ‘Taió’ were superior, combining high and stable productivity with low disease severity in both locations. Conversely, the cultivar ‘Ponteio’ showed high yield only in Lages, suffering a severe reduction and higher susceptibility in Agronômica. ‘Ceronte’ was the most susceptible genotype, reaching 86.25% severity in Agronômica. The Spearman correlation confirmed a negative and significant impact of blast on forage productivity, with a stronger correlation in Agronômica (ρ = −0.689) than in Lages (ρ = −0.497). These findings underscore that the GxA interaction is primarily driven by the differential resistance response to blast. ‘Altovale’ and ‘Taió’ are the most recommended options. It is concluded that the selection of ryegrass genotypes must be conducted in the target environment, considering blast as a primary selection factor to ensure the stability and sustainability of forage production in southern Brazil.

## Introduction

Annual ryegrass (*Lolium multiflorum* Lam.) is the main winter forage grass used in livestock production systems in subtropical climate regions, such as southern Brazil. Its high nutritional quality, palatability, and elevated productive potential make it a strategic resource for intensifying ruminant production, whether through direct grazing or as hay or silage. Additionally, its inclusion in integrated crop-livestock production systems contributes to soil protection and nutrient cycling, optimizing land use in rotation with summer crops (RECH et al., 2022).

However, the sustainability and productivity of ryegrass-based systems are constantly threatened by diseases, which can cause significant dry matter losses and reduce the forage’s nutritional value. Among the diseases that affect the crop, blast, caused by the fungus *Magnaporthe oryzae* (syn. *Pyricularia oryzae*), stands out as one of the main limiting factors, especially in years with favorable climatic conditions, such as high temperatures and high relative humidity (NUNES and MITTELMANN, 2009).

The development and utilization of annual ryegrass genotypes with genetic resistance constitute the most effective, economical, and environmentally sustainable management strategy for blast control. However, the expression of resistance, as well as the productive potential, is frequently influenced by the significant genotype-by-environment (GxE) interaction. A genotype with superior performance in a specific location or year may not maintain the same performance under different edaphoclimatic conditions or pathogen virulence spectra. This interaction requires breeding programs to evaluate their materials in a multi-environment trial network to identify genotypes with broad adaptability or specific adaptation to production niches (BORÉM et al., 2021).

In this context, regional breeding programs, such as the one Epagri conducts in Santa Catarina, Brazil, play a fundamental role in developing cultivars adapted to local conditions. Given the above, this study evaluated the productive performance and reaction to blast disease of commercial cultivars and advanced annual ryegrass lines in two edaphoclimatic environments in the state of Santa Catarina. The study hypothesized that the regionally developed genotypes would exhibit greater stability, productivity, and resistance to the disease, especially in the environment more favorable to the occurrence of blast.

## Materials and Methods

The experiment for evaluating annual ryegrass (*Lolium multiflorum*) cultivars and lines was conducted in Lages and Agronômica, Santa Catarina state, Brazil.

The experimental design was a randomized complete block design (RCBD) with four replications. The treatments consisted of eight annual ryegrass genotypes, namely six commercial cultivars (SCS316 CR Altovale, Ceronte, Feroz, SCS317 Centenário, Empasc 304, and BRS Ponteio) and two experimental lines (Taió and Eron).

The experimental plots consisted of 8 rows, 4.5 meters long, with 0.2 meters spacing between them, totaling an area of 7.2 m^2^ per plot. A spacing of 1 meter was maintained between the plots and between the blocks. Sowing was carried out using a density of 25 kg/ha of pure and viable seeds. Fertilization followed the technical recommendations for annual winter grasses, based on the soil analysis of the experimental area. Weed control was performed as needed.

Forage yield assessments were conducted by cutting the plants when they reached an average height of 25 cm, leaving a residue height of 10 cm. Two cuts were performed in each experimental plot, with each sample covering an area of 0.25 m^2^.

After evaluating the useful area, the remainder of the plot was equally lowered to 10 cm, and the plant material was removed from the site.

### The analyzed variables included

- **Dry mass yield**: The dry mass yield per hectare was estimated by summing the values of the two samples per plot, which provided the dry mass production in 0.5 m^2^. This value was then used to estimate the yield for 10000 m^2^ (per hectare). The sample collected in the field was weighed on a precision analytical balance after being placed in an oven at 65°C for approximately 72 hours. Once the sample reached a constant weight, the dry sample weight was obtained.
- **Disease tolerance**: Tolerance to disease attack was evaluated, with a specific focus on blast (*Pyricularia oryzae*). The severity of leaf blast was assessed by visual evaluation of the plot using a 0 to 100% scale, where 0 represents no lesions observed and 100 represents the total death of the plants in the plot.

### Statistical analysis

The data collected for the variables dry matter yield and blast severity were subjected to analysis of variance (ANOVA) according to the statistical model for a two-factor experiment (genotype × location). The statistical model used was:

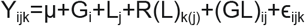

Where:

- Y_ijk_ is the observed value;
- μ is the overall mean;
- G_i_ is the effect of the i-th genotype;
- L_j_ is the effect of the j-th environment (or location);
- R(L)_k(j)_ is the effect of the k-th replication within the j-th environment;
- (GL)_ij_ is the interaction effect;
- ϵ_ijk_ is the random experimental error

The verification of the assumptions of the model’s homogeneity of variances and normality of residuals was performed using Levene’s and Shapiro-Wilk’s tests, respectively, both at a 5% significance level.

For the yield variable, the ANOVA assumptions were met, and the analysis was conducted with the original data.

For the blast severity variable, the assumptions of homogeneity of variances and normality of residuals were violated. Therefore, the data were transformed using the arc sine of the square root of the proportion function [y’ = asin(sqrt(y/100))] to stabilize the variance and approximate normality. ANOVA and subsequent mean comparison tests were performed on the transformed data.

For the main and interaction significant effects by the F-test, means were compared using Tukey’s test at 5% error probability. To facilitate interpretation, the results for the blast severity variable were presented in tables containing the original arithmetic means, accompanied by the statistical grouping letters obtained from the analysis of the transformed data. All statistical analyses were performed using the R software (R Core Team, 2025).

### Correlation Analysis

The relationship between dry matter yield and blast severity was quantified through a correlation analysis. First, the assumption of bivariate normality was evaluated. Due to the non-adherence of the blast severity data to the normal distribution, a non-parametric method was chosen.

The Spearman’s rank correlation coefficient (rho) was calculated to measure the strength and direction of the association between the two variables. The analysis was performed using the original data, without transformation. The null hypothesis of no correlation (ρ = 0) was tested, and the significance level adopted for the test was p < 0.05. The analyses and the creation of scatter plots were performed with the aid of the R software (R Core Team, 2025).

## Results and Discussion

The analysis of variance revealed a statistically significant interaction (p < 0.05) between the genotype and location factors for both evaluated variables: dry matter yield and blast severity. This result indicates that the ryegrass genotypes’ productive performance and disease reaction were inconsistent across the two environments, depending on the specific conditions of each location. Researchers frequently report this genotype x environment (GxE) interaction in the *Pyricularia oryzae* pathosystem in grasses, given the pathogen’s high variability and the strong climatic influence on the disease epidemic.

### Productive performance and adaptation of ryegrass genotypes

The breakdown of the interaction for dry matter yield (Table 1) allowed us to identify genotype groups with distinct adaptation. In the Lages environment (Cfb climate), the genotypes ‘Altovale,’ ‘Ponteio,’ ‘Taió,’ and ‘Feroz’ composed the superior group; they showed no statistical differences among themselves and achieved the highest yields. However, we observed a clear reclassification in Agronômica (Cfa climate). ‘Altovale’ and ‘Taió’ maintained their high performance, again ranking among the most productive. In contrast, the ‘Ponteio’ genotype suffered a drastic reduction in productivity; it moved from the elite group in Lages to the lowest-yielding group in Agronômica, becoming statistically similar to the ‘Ceronte’ cultivar.

**Table 1:** Analysis of the genotype by location interaction for dry matter yield (kg.ha-1). Means were compared using Tukey’s test at the 5% significance level. Means within a column followed by the same letter do not significantly differ.

| Gen | Lages | Agronômica |
| --- | --- | --- |
| Alto Vale | 4513.55 a | 4201.56 ab |
| Ponteio | 4347.62 a | 2504.23 e |
| Taió | 4263.99 a | 4149.71 abc |
| Feroz | 4183.72 ab | 3939.09 abcd |
| Eron | 3406.50 abcde | 2949.71 cde |
| Centenário | 3364.20 abcde | 3031.19 bcde |
| Empasc 304 | 3036.45 bcde | 3363.11 abcde |
| Ceronte | 2891.45 de | 2272.03 e |

The notable stability and high productivity of the ‘Altovale’ and ‘Taió’ genotypes in both locations, especially in Agronômica’s warmer and more humid environment, can be attributed to their origin. Having been selected in the Alto Vale do Itajaí, a region with a Cfa climate, these materials likely possess adaptive traits for higher temperature and humidity conditions, which favor the development of diseases like blast. Conversely, the ‘Ponteio’ genotype’s performance is a classic example of GxA interaction, demonstrating specific adaptation to Lages’ milder conditions and lower disease pressure (Cfb) and a clear lack of adaptation to the Agronômica environment.

### Reaction to blast and environmental Influence

The GxE interaction was even more evident for blast severity (Table 2). Disease pressure drastically increased in Agronômica compared to Lages. The ‘Ceronte’ genotype reached an average severity of 86.25% in Agronômica, revealing its extreme susceptibility, while in Lages, its severity was 22.75%; Nunes and Mittelmann (2017) reported similar results. This result corroborates the literature, which indicates that Cfa climate conditions (higher temperatures and elevated humidity) extremely favor *Pyricularia oryzae* epidemic progression.

**Table 2:** Unfolding of the genotypes x locations interaction for blast severity (%). Grouping letters were obtained using the Tukey test at 5% on transformed data (arc sine), but the means presented are the original values.

| Gen | Agronômica | Lages |
| --- | --- | --- |
| Ceronte | 86.25 a | 22.75 bc |
| Ponteio | 25.00 b | 21.00 bcd |
| Empasc 304 | 16.25 cdefg | 18.75 bcde |
| Centenário | 12.75 efgh | 17.25 bcdef |
| Alto Vale | 11.75 fgh | 14.50 defgh |
| Eron | 10.50 fgh | 13.50 efgh |
| Taió | 9.75 h | 12.50 efgh |
| Feroz | 10.00 gh | 10.25 gh |

The genotypes ‘Feroz’, ‘Taió’, and ‘Altovale’ stood out because they presented the lowest disease severity levels in both locations, indicating a broad and stable resistance profile. The resistance expressed by ‘Taió’ and ‘Altovale’ reinforces the hypothesis that their selection in the Alto Vale do Itajaí environment (Cfa) resulted in the incorporation of effective resistance genes against pathogen populations prevalent under high inoculum pressure conditions.

### Impact of the disease on yield

Spearman’s correlation analysis (Table 3) confirmed the significant negative impact of blast on ryegrass forage productivity. We observed a negative correlation in both locations, which was stronger in Agronômica (ρ = −0.689; p < 0.001) than in Lages (ρ = −0.497; p < 0.01).

**Table 3:** Spearman’s correlation coefficients between blast severity and dry matter yield, by location, Santa Catarina, Brazil, 2024.

| Local | Coefficiente de Spearman (rho) | Valor-p | Significância |
| --- | --- | --- | --- |
| Agronômica | -0.689 | 0.0000 | *** |
| Lages | -0.497 | 0.0038 | ** |

This difference in the strength of the correlation provides a relevant insight. In Agronômica, where the disease manifested severely and with high variability among genotypes, blast became a primary factor limiting yield. For example, the decrease in productivity of the cultivar ‘Ponteio’ in Agronômica, which had greater susceptibility (25% severity), compared to the more resistant genotypes (Figure 1), can be directly attributed to the blast, confirming the study conducted by Cruz et al. (2025). In Lages, although the disease still negatively impacted production, the lower pathogen pressure meant other genetic and environmental factors had a relatively greater influence in determining the final yield.

**Figure 1.**
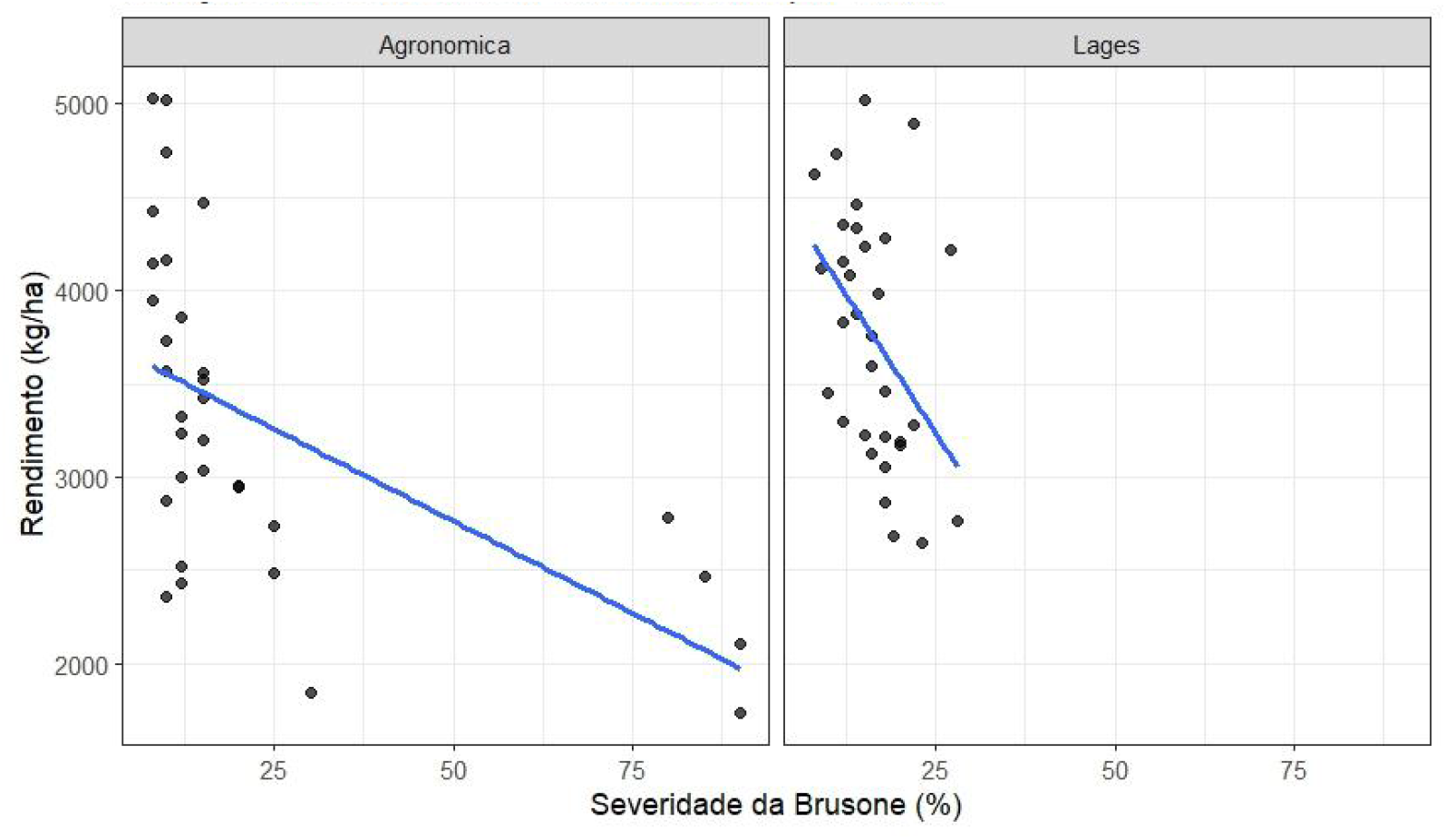
Influence of blast on dry matter yield of different annual ryegrass genotypes by location, Santa Catarina, Brazil, 2024.

## Conclusions

The results demonstrate the existence of a strong genotype x environment interaction, mainly governed by the differential response of the genotypes to blast disease under distinct climatic conditions. The ‘Altovale’ and ‘Taió’ genotypes, originating from a high disease pressure environment (Cfa), stood out as superior because they combined high productivity and stable resistance, making them the most recommended options for both regions. The ‘BRS Ponteio’ cultivar demonstrated good adaptation only to the lower disease pressure environment (Cfb), while ‘Ceronte’ proved highly susceptible and low-yielding, which disqualifies it for regions with a history of blast. Therefore, annual ryegrass genotype selection must occur in the target environment, considering blast as a main selection factor to ensure the stability and sustainability of forage production.

